# The hidden diversity of ancient bornaviral sequences from X and P genes in vertebrate genomes

**DOI:** 10.1101/2022.05.06.490909

**Authors:** Bea Clarise B. Garcia, Yahiro Mukai, Keizo Tomonaga, Masayuki Horie

## Abstract

Endogenous bornavirus-like elements (EBLs) are heritable sequences derived from bornaviruses in vertebrate genomes that originate from transcripts of ancient bornaviruses. EBLs have been detected using sequence similarity searches such as tBLASTn, whose technical limitations may hinder the detection of EBLs derived from small and/or rapidly evolving viral X and P genes. Indeed, no EBLs derived from the X and P genes of orthobornaviruses have been detected to date in vertebrate genomes. Here, we aimed to develop a novel strategy to detect such “hidden” EBLs. To this aim, we focused on the 1.9 kb read-through transcript of orthobornaviruses, which encodes a well-conserved N gene and small and rapidly evolving X and P genes. We show a series of evidence supporting the existence of EBLs derived from orthobornaviral X and P genes (EBLX/Ps) in mammalian genomes. Furthermore, we found that an EBLX/P is expressed as a fusion transcript with the cellular gene, *ZNF451*, which potentially encodes the ZNF451/EBLP fusion protein in miniopterid bat cells. This study contributes to a deeper understanding of ancient bornaviruses and co-evolution between bornaviruses and their hosts. Furthermore, our data suggest that endogenous viral elements detected thus far are just the tip of the iceberg, and further studies are required to understand ancient viruses more accurately.

## Introduction

Ancient viral sequences can be found within the genomes of organisms in the form of endogenous viral elements (EVEs) (1). EVEs are derived from the integration of viral sequences into the host germline DNA and transfer of the integrated sequences to subsequent progenies (1, 2). As molecular fossils of ancient viruses, EVEs have expanded our knowledge of the diversity, host range, and long-term evolution of viruses (1). Furthermore, some EVEs are co-opted by hosts to function as novel gene regulatory elements, non-coding RNAs, and proteins (2).

Most EVEs are derived from retroviruses, as the viruses reverse-transcribe and integrate the viral genomes into the host genomes using virally encoded enzymes. However, non-retroviral EVEs have also been identified. Endogenous bornavirus-like elements (EBLs) are EVEs derived from bornaviruses (family *Bornaviridae*), a group of non-segmented negative-strand RNA viruses (3, 4). Although bornaviruses are RNA viruses, evidence strongly suggests that bornaviral mRNA is reverse transcribed and integrated into the host genome via the activity of the long interspersed nuclear element 1 (LINE-1) retrotransposon (3, 5). We previously revealed an abundance of EBLs in vertebrate genomes derived from the bornaviral genes N, P, M, G, and L (S1 Fig), which are referred to as EBLN, EBLP, EBLM, EBLG, and EBLL, respectively (6). These EBLs originated from ancient bornaviruses of all genera in the family *Bornaviridae*: *Orthobornavirus, Carbovirus*, and *Cultervirus*.

Despite the rigorous search, there has been a bias in the detection of EBLs. EBLNs comprise the majority of the detected EBLs, while no EBLs derived from the X and P genes have been reported except for a few carboviral EBLPs (3, 6-10). We proposed two hypotheses to explain the lack of EBLX and EBLP. First, X/P mRNA, which encodes the X and P proteins, may have a low propensity for reverse-transcription and/or integration into the host genome. However, previous studies of Borna disease virus 1 (BoDV-1; genus *Orthobornavirus*) showed that X/P mRNA is the most abundant transcript among the viral mRNAs (11, 12), which should therefore provide enough chances for integration. Moreover, we have previously demonstrated that the X/P mRNA of BoDV-1 can be reverse-transcribed and integrated into the genome of host cells (3, 7). This led us to the second hypothesis, which states that EBLs derived from the X and P genes are present but are difficult to detect due to technical limitations of the sequence similarity search. EBLs have been routinely searched using tBLASTn; however, this can be an unsuitable method for detecting small and rapidly evolving genes (8). Indeed, the X and P genes are small and the least conserved among bornaviral genes (Fig 1). If the second hypothesis is correct, some EBLs may remain undetected. Therefore, we could be missing important insights into ancient bornaviruses and the co-evolution of bornaviruses and their hosts.

**Figure 1.**
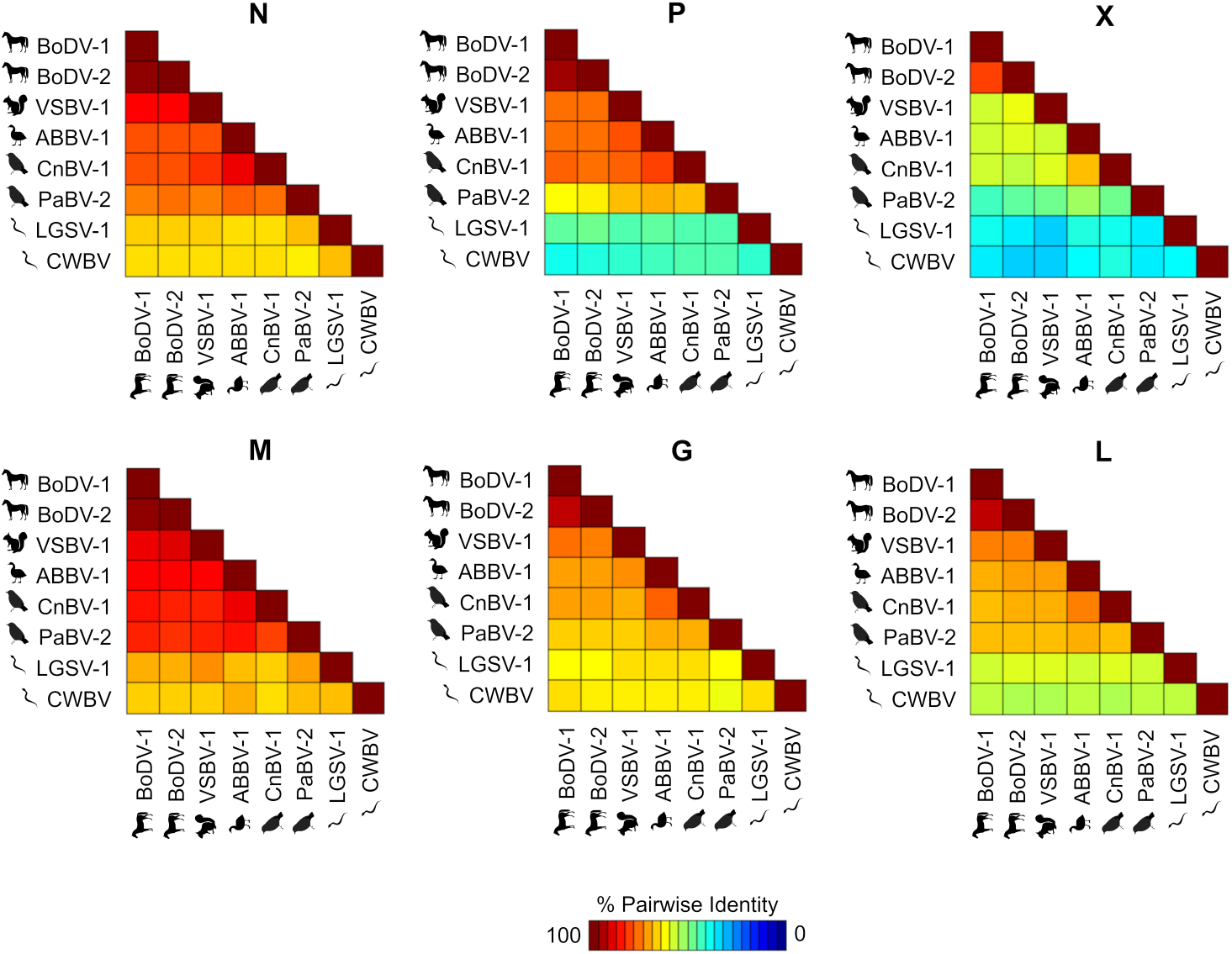
Amino acid identities between extant orthobornaviral proteins. Pairwise identities between the amino acid sequences of extant orthobornaviral proteins was analyzed using Sequence Demarcation Tool (35). The animal silhouettes show the representative hosts of viruses.

In this study, we sought to develop a novel strategy to detect these hidden EBLs by focusing on the nature of the read-through transcription of orthobornaviruses that produce a transcript encoding a highly conserved N gene and rapidly evolving X and P genes. We show evidence supporting the existence of several EBLX/Ps in vertebrate genomes. Furthermore, an EBLX/P in miniopterid bats appear to be transcribed as chimeric transcripts with a host gene, suggesting a potential co-option of EBLs by the host.

## Results

### Miniopterid bat EBLN is possibly derived from an N/X/P transcript

We previously identified miEBLN-1, an EBLN derived from an ancient orthobornavirus that retains an intact N ORF in miniopterid bats (13). The miEBLN-1 locus is present in the genomes of miniopterid bats, specifically in those of *Miniopterus natalensis, M. schreibersii*, and *M. fuliginosus*, but is empty (i.e., syntenic loci lacking miEBLN-1) in the genomes of *Rhinolophus ferrumequinum* and *Phyllostomus hastatus*. We re-examined the presence of miEBLN-1 using the current genome datasets and found the miEBLN-1-empty locus in *Tadarida brasiliensis* (Fig 2a and S2a Fig).

**Figure 2.**
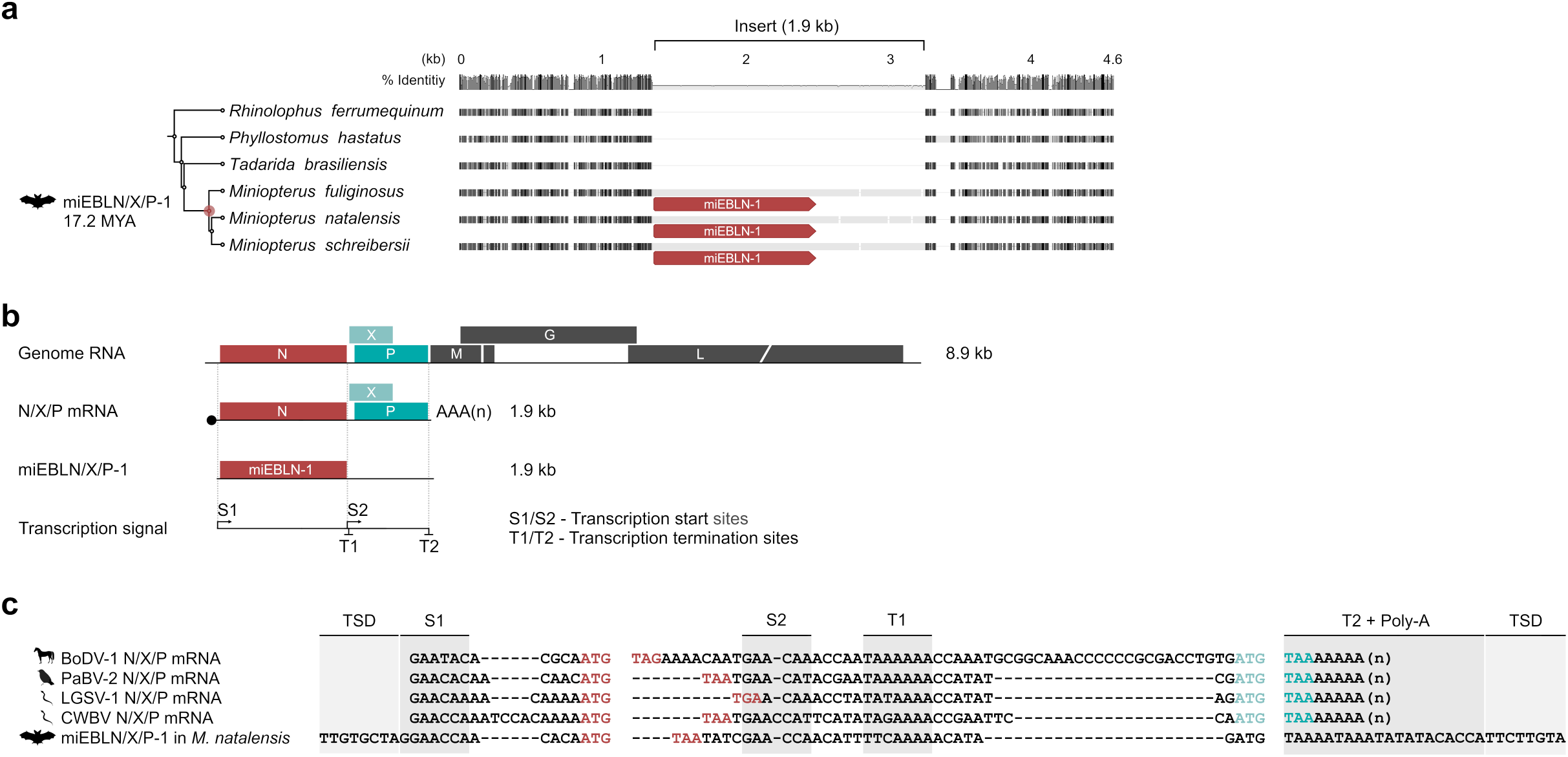
Characterization of miEBLN-1 as an EBL derived from N/X/P transcript. **(a)** Gene orthology analysis of the miEBLN/X/P-1 locus. Multiple alignment of the deduced syntenic loci between the indicated species is shown. Black and gray boxes show aligned regions. Thin gray lines indicate alignment gaps. Species phylogeny and minimum integration age of miEBLN/X/P-1 are indicated on the left based on TimeTree (38). The full version of this alignment is available in S1 Fig. **(b)** Schematic diagram of orthobornaviral genomic RNA, 1.9-kb N/X/P mRNA, and miEBLN/X/P-1. Transcription signals (S1, S2, T1, and T2) are shown. **(c)** Nucleotide sequence alignment of extant orthobornaviral N/X/P mRNA and miEBLN/X/P-1. Red letters indicate the start and stop codons of the N gene. Light and dark teal blue letters indicate the start codon of X gene and the stop codon of P gene, respectively. TSD, target site duplication. The full version of this alignment is available in S2 Fig.

Based on the nucleotide alignment, we deduced the insertion size of the miEBLN-1 locus to be 1.9 kb, consisting of a 1.2-kb miEBLN-1 region and a 0.7-kb unannotated region. Therefore, we carried out a series of analyses to understand the source of the 0.7-kb unannotated region. First, we performed a BLASTn search (14) against the NCBI nt database (15) using the nucleotide sequence of the 0.7-kb unannotated region as a query. The hits consisted only of miEBLN-1 sequences (data not shown). Next, we assessed whether the 0.7-kb unannotated region is derived from a short interspersed nuclear element (SINE) due to the precedence of chimeric integration comprising nucleotide sequences from an N transcript and SINE (3, 5). We performed a BLASTn search against the miniopterid bat genomes using the nucleotide sequence of the 0.7-kb region as a query. We only detected five BLAST hits in the genomes of *M. natalensis* and *M. schreibersii*, which we identified as EBLs in the later sections. We also analyzed the sequence of the 0.7-kb region by CENSOR (16), but no hits were detected. These results indicated that the 0.7-kb fragment is not derived from a SINE.

We noticed that the insertion, including miEBLN-1, is almost similar in size to the N/X/P mRNA of extant orthobornaviruses (Fig 2a and S2a Fig), which led us to speculate that the insertion may be derived from an N/X/P transcript. N/X/P mRNA is produced by read-through of the termination signal T1, resulting in a 1.9 kb mRNA containing the transcription signal sequences S1, T1/S2, and T2 (Fig 2b) (12). Therefore, we investigated the presence of these signal-like sequences in the 1.9-kb miEBLN-1 insertion. We found transcription signal-like sequences at reasonable positions with respect to extant N/X/P transcripts (Fig 2c and S3b Fig). A poly-A stretch was also observed immediately downstream of the T2 signal, indicating that the insertion originated from an mRNA transcript (Fig 2c and S3b Fig). Furthermore, a tandem duplication sequence, TTGTGCTA/TTCTTGTA, was observed immediately upstream and downstream of S1 and T2-poly-A sequences, respectively (Fig 2c and S3b Fig). This is a potential target site duplication (TSD), which is a hallmark of LINE-1-mediated mRNA integration (17). These results suggested that the 1.9-kb insertion, including miEBLN-1, is derived from an ancient N/X/P transcript integrated by LINE-1. Therefore, we tentatively refer to this locus as miEBLN/X/P-1.

### Additional EBLs possibly derived from N/X/P transcripts in miniopterid bat genomes

As described above, orthobornaviruses express mRNAs encoding N (1.2 kb), X and P (X/P mRNA, 0.7 kb), or N, X, and P (N/X/P mRNA, 1.9 kb) (Fig 2a). If miEBLN/X/P-1 is truly derived from N/X/P mRNA, there may be additional EBLs derived from N/X/P or X/P mRNA. To examine this possibility, we conducted a BLASTn search against bat genomes using the nucleotide sequence of miEBLN/X/P-1 as a query. We obtained six hits from the genomes of *M. natalensis* and *M. schreibersii* (Fig 3a and S1 Table). While four hits were aligned in the entire N/X/P region of miEBLN/X/P-1 (Fig 3a, red lines), interestingly, two other hits were aligned only to the putative X/P region (Fig 3a blue lines). Note that the N regions of four hits aligned to entire miEBLN/X/P-1 were detected as EBLNs in a previous study (6).

**Figure 3.**
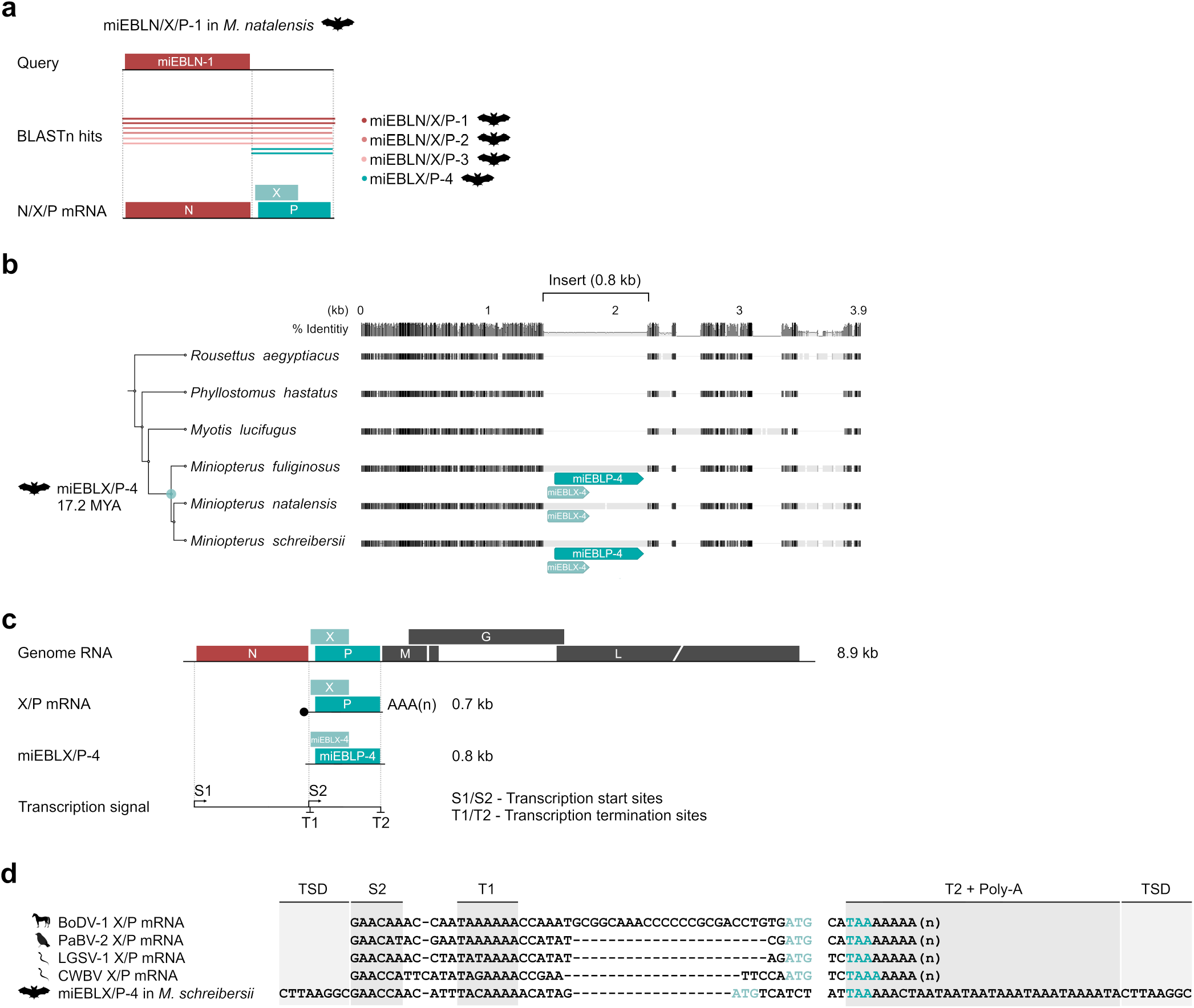
Identification of an EBL derived from X/P transcript in miniopterid bats. **(a)** Schematic diagram of the BLASTn search against the bat whole genome shotgun sequence database, using miEBLN/X/P-1 as a query. The BLAST hits are shown by dark to light read or teal blue lines corresponding to the EBLs indicated on the right. **(b)** Gene orthology analysis of the miEBLX/P-4 locus. Species phylogeny and deduced minimum integration age of miEBLX/P-4 are indicated on the left based on TimeTree (38). miEBLX-4 and miEBLP-4 ORFs are shown in light and dark teal blue, respectively. The full version of this alignment is available in S3 Fig. **(c)** Schematic diagram of extant orthobornaviral genomic RNA, X/P mRNA, and miEBLX/P-4. Transcription signals (S2, T1, and T2) are indicated. **(d)** Nucleotide sequence alignment of extant orthobornaviral X/P transcripts and miEBLX/P-4. Light and dark teal blue letters indicate the start codon of X gene and the stop codon of P gene, respectively. TSD, target site duplication. The full version of this alignment is available in S4 Fig.

We first assessed hits that aligned to the entire region of miEBLN/X/P-1. Four hits showed 1.9 kb alignment lengths and high sequence similarities (76–84% nucleotide identity) to miEBLN/X/P-1 (S1 Table). The flanking regions of each pair of hit sequences were well aligned between *M. natalensis* and *M. schreibersii*, indicating that they are orthologous between these species (S7 Table). We tentatively refer to these loci as miEBLN/X/P-2 and miEBLN/X/P-3. We also confirmed the presence of orthologs of these EBLN/X/Ps in the genome of *M. fuliginosus* by PCR and Sanger sequencing of genomic DNA isolated from *M. fuliginosus* (S2b Fig and S7 Table).

Next, we examined the presence of miEBLN/X/P-2 or miEBLN/X/P-3-empty loci by comparative genomics. We observed that the miEBLN/X/P-3 empty locus is present in genomes of non-miniopterid bats such as *Tadarida brasiliensis* (S2b Fig and S7 Table). We did not obtain an alignment of the flanking region of miEBLN/X/P-2 with the other bat genomes and therefore could not identify its empty locus.

We then analyzed the nucleotide sequences of miEBLN/X/P-2 and miEBLN/X/P-3. Similar to miEBLN/X/P-1, both EBLN/X/Ps contained transcription signal-like sequences, and poly-A stretches were observed downstream of T2-like sequences (S3b Fig). Furthermore, tandem duplication sequences, TGTTCA/[T/C]GTTCA and GAGGAACG/GAGGGATG, were also observed immediately upstream of S1 and downstream of T2/poly-A in miEBLN/X/P-2 and miEBLN/X/P-3, respectively (S3b Fig). These results suggest that miEBLN/X/P-2 and miEBLN/X/P-3 are also derived from ancient N/X/P transcripts and were integrated into the genome of a miniopterid bat ancestor through LINE-1 activity.

### EBLs derived from X/P-like transcripts exist in miniopterid bat genomes

Next, we assessed the hits that aligned to the 0.7-kb putative X/P region of miEBLN/X/P-1 (Fig 2a, teal blue lines). Two hits from *M. natalensis* and *M. schreibersii* showed 0.8 kb alignment lengths and 75% nucleotide identity to the 0.7-kb X/P region (S1 Table). The flanking regions of the hit sequences were well aligned, indicating that the loci are orthologous between these species (Fig 3b and S4a Fig; S7 Table). We also found an orthologous locus in the genome of *M. fuliginosus* using PCR and Sanger sequencing (Fig 3b and S4a Fig; S7 Table). Furthermore, we detected empty loci in genomes of non-miniopterid bats, genomes such as *Myotis lucifugus* (Fig 3b and S4a Fig; S7 Table).

The orthobornaviral X/P mRNA contains the transcription signal sequences S2/T1 in the 5’-UTR and T2 in the 3’-UTR (Fig 3c). We observed that S2/T1 and T2-like sequences are present in the 0.7-kb insertion, which are located at positions similar to those in extant X/P mRNA (Fig 3d and S5b Fig). Poly-A stretches were also observed downstream of the T2-like sequences. Furthermore, the S2 and poly-A stretches were flanked by TSDs, CTTAAGGC/CTTAAGGC (Fig 3d and S5b Fig). Taken together, these results indicated that the insertion may have been derived from an ancient X/P transcript and integrated into the genome of a miniopterid bat ancestor through LINE-1 activity. We tentatively refer to the insertion as the miEBLX/P-4.

### Preservation of intact ORFs in miEBLX/P-4 and characterization of the putatively encoded proteins

Most EBLs do not retain viral ORFs because of the accumulation of mutations after integration. Remarkably, miEBLX/P-4 retains intact X- and P-like ORFs in different reading frames, which is consistent with the extant orthobornaviral X/P mRNA (Fig 3c). We named these ORFs as miEBLX-4 and miEBLP-4. Notably, miEBLP-4 in *M. natalensis* is disrupted by a premature stop codon.

Although there is very low sequence similarity between the amino acid sequences of these ORFs and extant orthobornaviral proteins, further analyses showed that miEBLX-4 and miEBLP-4 are intrinsically similar to extant orthobornaviral X and P proteins.

First, we examined the miEBLX-4 ORF. The ORF consists of 109 codons, which is longer than that of extant orthobornaviral X (86–90 codons). We then predicted the secondary structure and analyzed the hydrophobicity of the putative miEBLX-4 protein with respect to extant X proteins. Both extant X and miEBLX-4 were predicted to contain an alpha helix in the N-terminal region (Fig 4a and S6a Fig). The hydropathy plot showed similar hydrophobicity patterns for these proteins (S7a Fig).

**Figure 4.**
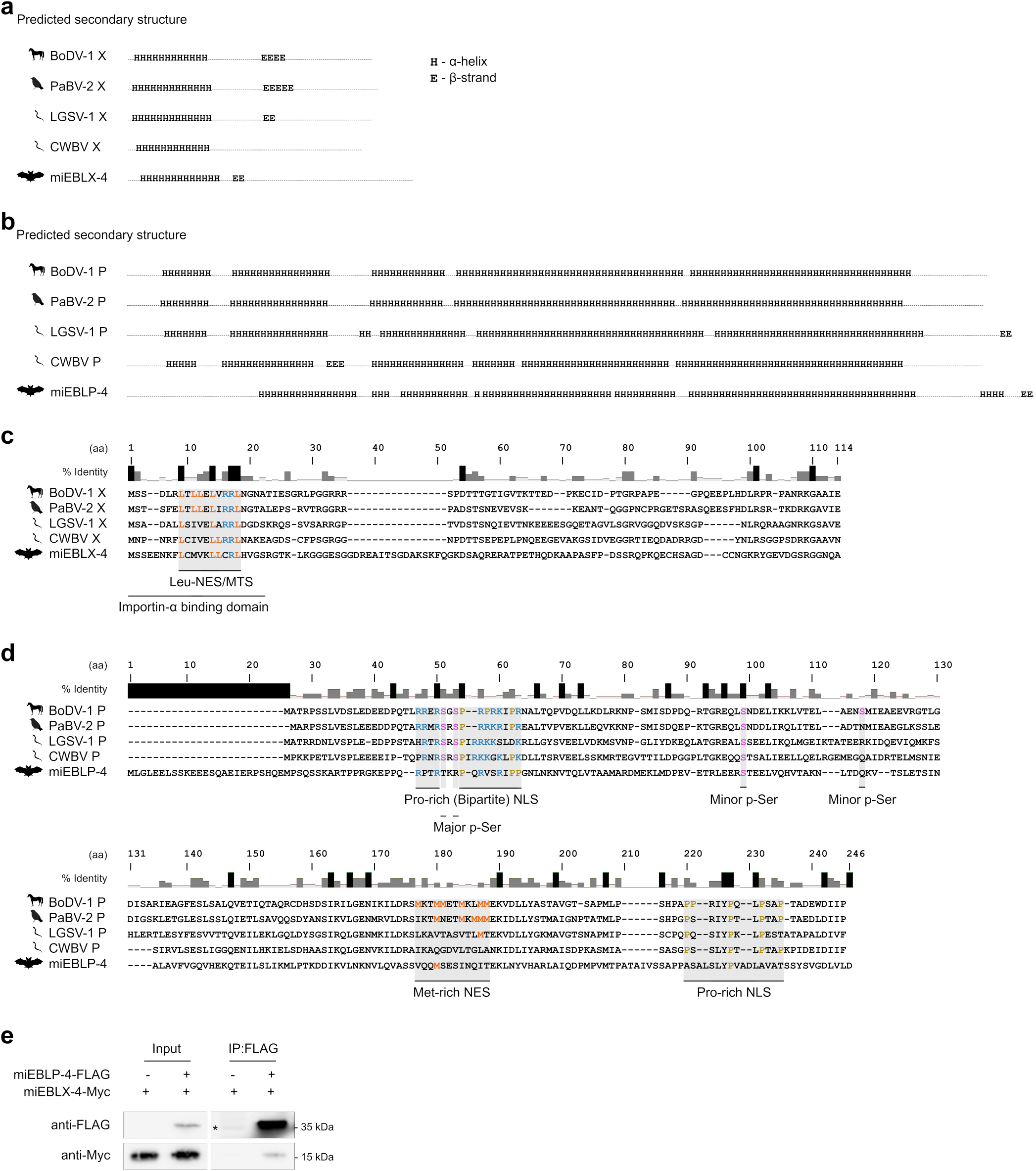
Conservation of properties between putative EBLX and EBLP and viral X and P proteins. **(a** and **b)** Predicted secondary structure of extant orthobornaviral X and miEBLX-4 proteins **(a)**, and extant orthobornaviral P and miEBLP-4 proteins **(b)**. H, α-helix. E, β-strand. The data including amino acid residues are shown in Figs. S5 and S6. **(c** and **d)** Amino acid sequence alignment of extant orthobornaviral X and miEBLX proteins **(c)**, and extant orthobornaviral P and miEBLP-4 proteins **(d)**. Shaded areas indicate the functional domains previously characterized in BoDV-1 X and P proteins. The conserved amino acid residues important for the functions of viral X or P protein are highlighted in colors. NLS, nuclear localization signal; NES nuclear export signal; p-Ser, phosphorylated serine residue. **(e)** Co-immunoprecipitation of Myc-tagged miEBLX-4 with FLAG-tagged miEBLP-4. FLAG-tagged miEBLP-4 was immunoprecipitated, and then analyzed by western blotting using the indicated antibodies. *, Non-specific band.

We also determined whether the functional domains are conserved between extant X and miEBLX-4. Amino acid sequence alignment showed that the N-terminal region is relatively well conserved and houses functional domains previously characterized in BoDV-1 X (Fig 4c). This region includes the leucine-rich domain functioning both as a CRM1-dependent nuclear export signal (18) and a mitochondria-targeting signal (19), and is overlapped by the importin-alpha binding domain (20). The alignment also showed that leucine and arginine residues that are critical in the importin-alpha-binding domain (Fig 4c; orange and blue, respectively) are conserved among extant X and miEBLX-4.

Next, we examined the miEBLP-4 ORF. The ORF consists of 233 codons, longer than those of extant orthobornaviral P (201–208 codons). The predicted secondary structures and hydrophobicity pattern of miEBLP-4p were similar to those of the modern counterparts (Fig 4d, S6b Fig, and S7b Fig). Alignment of the amino acid sequences revealed that some of the functional domains previously characterized in BoDV-1 P may be shared between extant P and miEBLP-4 (Fig 4d). Basic and proline residues important for the nuclear localization signal (NLS) activity (Fig 4d, blue) (21, 22) in the N-terminal NLS were prominent in both proteins.

We also explored the interaction between miEBLX-4 and miEBLP-4, which is a notable characteristic of the orthobornaviral X and P proteins (23). We expressed Myc-tagged miEBLX-4 (miEBLX-4-Myc) and FLAG-tagged miEBLP-4 (miEBLP-4-FLAG) in 293T cells and then immunoprecipitated them using an anti-FLAG antibody. miEBLX-4-Myc was co-immunoprecipitated with miEBLP-4-FLAG, indicating the interaction between these proteins (Fig 4e).

These results showed that miEBLX-4 and miEBLP-4 share similar intrinsic properties with the extant X and P proteins. Furthermore, together with the above data, they strongly suggest that miEBLN/X/P-1-3 and miEBLX/P-4 are indeed derived from ancient viral N/X/P and X/P mRNAs, respectively.

### Diverse EBLX/Ps are found in vertebrate genomes

To further identify the presence of other EBLX/Ps, we conducted TFASTX and TFASTY searches against vertebrate genomes using the amino acid sequences of miEBLX-4 and miEBLP-4 as queries. TFASTX and TFASTY are sequence similarity search programs that allow frameshifts for alignment (24). As a result, 65 unique hits (S2-5 Tables) were detected in the genomes of mammals (class Mammalia), reptiles (class Reptilia), amphibians (class Amphibia), and ray-finned fish (class Actinopterygii). The majority of these hits were detected in the bat genomes, including miEBLN/X/P-1, miEBLN/X/P-2, miEBLN/X/P-3, and miEBLX/P-4.

Comparative genomic analyses strongly suggest that at least some of the TFASTX/Y hits are indeed derived from insertions of ancient viral X/P transcripts, as follows. TFASTX/Y hits in the bat families Mormoopidae and Phyllostomidae (mpEBLX/P) are orthologous, and their empty loci are present in other lineages of bats, such as *Noctilio leporinus* (Fig 5a and S4b Fig). For a TFASTX/Y hit in the lone bat species *Craseonycteris thonglongyai* (ctEBLX/P), we could not find its ortholog, but its empty sites exist in other bat species (Fig 5b and S4c Fig). We further identified orthologous TFASTX/Y hits in the rodent superfamily Octodontoidea and their empty loci in other lineages of rodents (ocEBLX/P; Fig 5c and S4d Fig). We also identified transcription signal-like sequences and TSDs within and flanking the above TFASTX/Y hits, respectively (Fig 5d and S5b Fig). These results strongly suggested that at least some of the sequences detected by TFASTX and TFASTY are bona fide EBLX/Ps.

**Figure 5.**
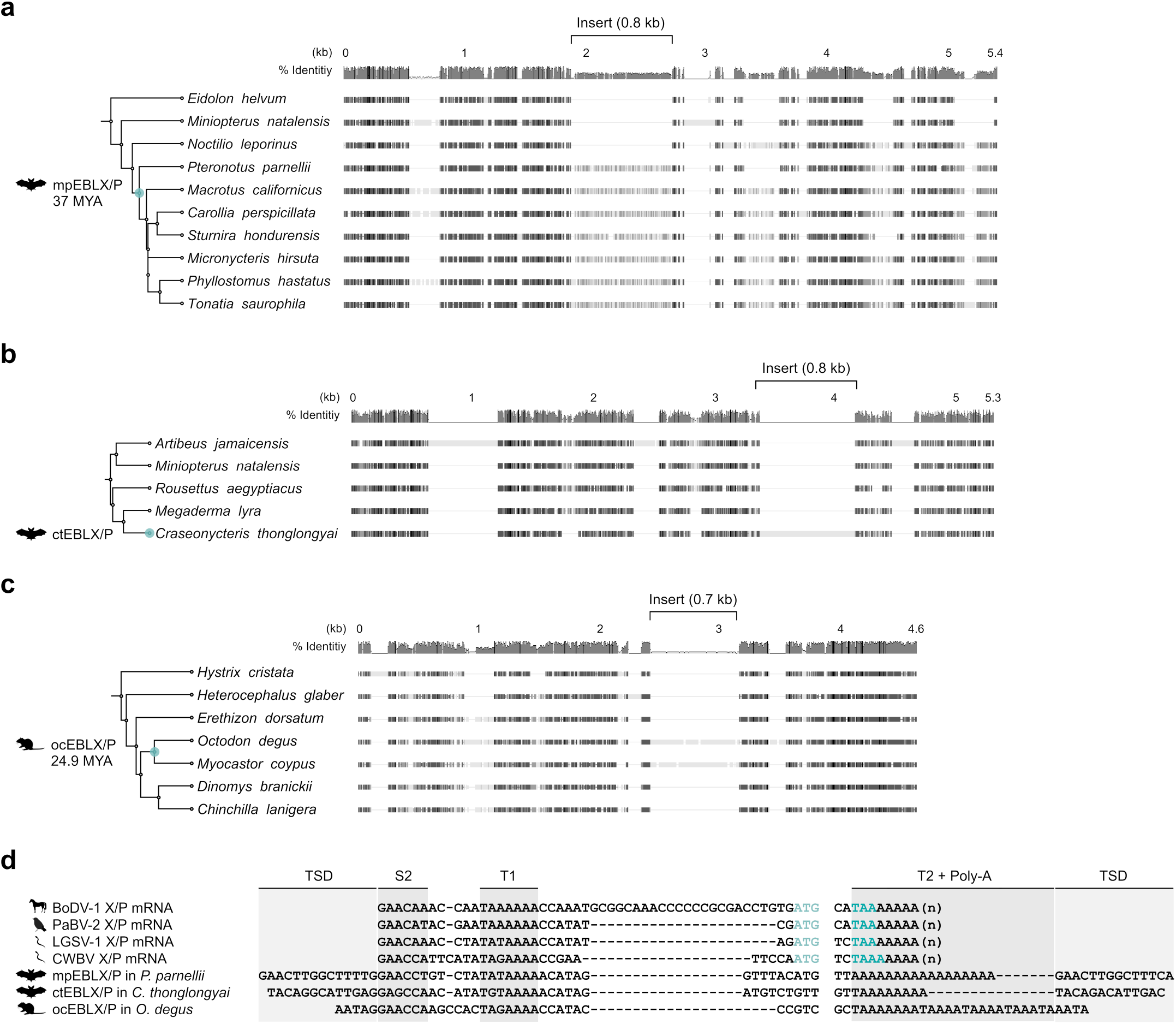
Identification of diverse EBLX/Ps in the vertebrate genomes. **(a-c)** Gene orthology analysis of each EBLX/P locus. Species phylogeny and deduced minimum integration age of each EBLX/P are indicated on the left based on TimeTree (38). The full version of this alignment is available in S3 Fig. **(d)** Nucleotide sequence alignment of extant orthobornaviral X/P transcripts and EBLX/Ps. Black and gray boxes show aligned regions. Thin gray lines indicate alignment gaps. Light and dark teal blue letters indicate the start codon of X gene and the stop codon of P gene, respectively. TSD, target site duplication. The full version of this alignment is available in S4 Fig.

### Possible co-option of miEBLX/P-4 in miniopterid bats

Some EVEs, including EBLs, retain intact ORFs which encode proteins that play roles in host-related functions (25). As described above, miEBLX/P-4 contains intact ORFs corresponding to orthobornaviral X and P genes. Therefore, we analyzed miEBLX/P-4 in detail.

We analyzed the expression of miEBLX/P-4 by mRNA sequencing of *M. schreibersii*-derived SuBK12-08 cells. miEBLX/P-4 is located within the intron region of *ZNF451* gene. The read coverage of the miEBLX/P-4 locus was comparable to that of *ZNF451* (Fig 6a). Interestingly, the mapping pattern revealed the existence of an alternative splicing site for *ZNF451* that could produce a chimeric transcript comprising ZNF451 exons 1–4 and the miEBLX/P-4 locus (Fig 6a).

**Figure 6.**
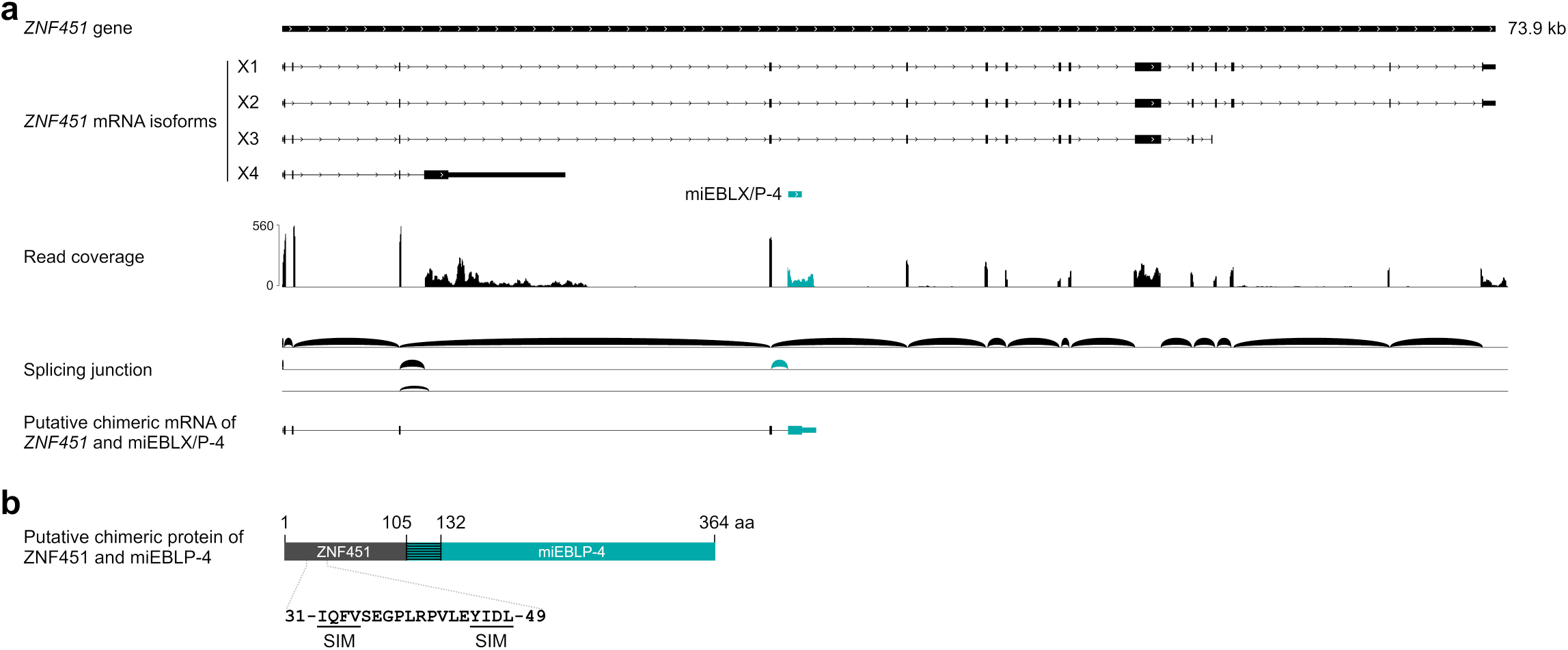
Alternative splicing generates *ZNF451*/*miEBLX/P-4* chimeric transcript. **(a)** Coverage and splicing junctions of mapped RNA reads from SuBK 12-08 cells (*M. schreibersii*) at the *ZNF451* locus where miEBLX/P-4 is located. The miEBLX/P-4 region, mapped read coverage graph on miEBLX/P-4, and splicing junction between ZNF451 and miEBLX/P-4 are shown in teal blue. **(b)** Schematic figure of putative chimeric ZNF451/miEBLP-4 protein. Shaded teal blue box SIM, SUMO interacting motif.

The putative chimeric transcript contains two ORFs consisting of more than 100 codons in different reading frames: a chimeric ORF of ZNF451 in-frame with miEBLP-4 or that of the miEBLX-4 (Fig 6b). The chimeric ORF potentially encodes a fusion protein comprising the N-terminal region of ZNF451, which contains a SUMO-interacting motif (SIM) domain, and miEBLP-4 (Fig 6b).

These results suggest that miEBLX/P-4 has been co-opted by miniopterid bats as a chimeric gene with *ZNF451*.

## Discussion

We previously demonstrated that the X/P mRNA of a modern orthobornavirus can be reverse-transcribed and integrated into the host genome (3, 7); however, no EBLX and only a few EBLPs have been reported (6). We hypothesized that the technical limitations of the sequence similarity search hinder the detection of EBLs derived from the X and P genes. EBLs have been routinely searched using tBLASTn, but this can be an unsuitable method for detecting small and rapidly evolving genes such as the X and P genes (8). In this study, our aim was to develop a novel strategy to detect these hidden EBLs. To this aim, we focused on the 1.9-kb read-through transcript of orthobornaviruses, consisting of a well-conserved N gene and small and rapidly evolving X and P genes (12), to detect hidden EBLX/Ps. We show a series of evidence implying that EBLN/X/Ps and EBLX/Ps are present in vertebrate genomes but are not detectable by BLAST searches (Figs 2 and 3). Interestingly, miEBLX/P-4 has maintained intact ORFs for more than 17 million years (Fig 3) and is expressed as a fusion transcript with the cellular gene, *ZNF451*, which potentially encodes the ZNF451/miEBLP-4 fusion protein (Fig 6). Therefore, this study revealed the presence of hidden EBLs, which could contribute to a deeper understanding of ancient bornaviruses and the co-evolution between bornaviruses and their hosts.

Here, we uncovered the potential co-option of miEBLX/P-4 by miniopterid bats, as indicated by the expression of the ZNF451/miEBLP-4 chimeric transcript, which potentially encodes a fusion protein consisting of the N-terminal region of ZNF451 and miEBLP-4 (Fig 6). To the best of our knowledge, this is the first report of a non-retroviral EVE being expressed as a chimeric transcript with a known host gene. Although we could not confirm the expression of the ZNF451/miEBLP-4 fusion protein due to the lack of antibodies, our data suggest that miEBLX/P-4 has been co-opted by miniopterid bats. Further studies are needed to understand the biological significance of miEBLX/P-4.

Our data strongly suggest that the genome organization and expression patterns of the N, X, and P genes have not substantially changed over the past million years. Here, we show that T1-read-through transcription occurred in ancient orthobornaviruses present at least 17.3 MYA (Fig 2). Furthermore, the ORF positions of the X and P genes of ancient orthobornaviruses were highly similar to those of modern orthobornaviruses (Fig 3). In addition, the sequences and order of transcription signals are highly conserved among ancient and modern orthobornaviruses (Figs 2 and 3). In BoDV-1, read-through transcription has been suggested to be involved in the regulation of viral gene expression (12). This hypothetical gene regulation mechanism has been maintained for more than 17.3 million years. Interestingly, our previous study showed that the order of S2-T1 is also conserved in some EBLNs in primates, which were estimated to be integrated into the host genomes more than 43 MYA (3, 5, 9). Thus, this gene regulatory mechanism may have been employed by orthobornaviruses earlier than 17.3 MYA.

The fundamental functions of viral X and P proteins may also be evolutionarily conserved between ancient and modern orthobornaviruses. We observed similarities in the intrinsic properties of extant orthobornaviral X and P proteins with miEBLX-4 and miEBLP-4 proteins despite their very low amino acid identities (Fig 4, S6 Fig, and S7 Fig). Furthermore, the interaction between X and P may have been conserved in ancient orthobornaviruses (Fig 4e). Therefore, although the evolutionary rates of X and P genes are high, the fundamental features and functions of X and P proteins may have been maintained for at least 17.3 million years.

For the EVE search, TFASTX/Y may be more suitable than tBLASTn, a conventional sequence similarity search for the detection of EVEs. After integration into the host genome, many EVEs have acquired random mutations, which caused frameshifts in the ORFs. Such frameshifted EVEs are a kind of “fragmented” EVEs and are thus less detectable by tBLASTn. In contrast, TFASTX/Y takes frameshifts into account (24) and can thus detect such fragmented EVEs more sensitively. We are currently conducting a benchmark analysis of tBLASTn and TFASTX/Y in detail, showing that some of the EBLX/Ps detected by TFASTX/Y cannot be detected by tBLASTn or other BLAST programs (data not shown). Therefore, TFASTX/Y can detect fragmented EVEs more sensitively than tBLASTn, which may contribute to identifying other hidden EVEs.

The EBLX/Ps identified in this study would represent only the tip of the iceberg on the hidden EVEs that could exist in the genome. In this study, the majority of EBLX/Ps were detected in bat and rodent genomes. Note that we failed to detect any EBLX/Ps in primate genomes, which contain numerous EBLs. Considering the number of EBLs in the genomes of primates (i.e., 24 orthobornaviral EBLs were detected in the human genome (6)), it is plausible that hidden EBLX/Ps exist in primates that are undetectable because of the inherent low amino acid similarities in X and P.

We may be able to detect such hidden EBLX/Ps by identifying other EBLN/X/P integrations. However, searching for other EBLN/X/Ps by manual curation is time-consuming. Thus, the development of an efficient searching method to identify approximately 1.9-kb insertions with EBLN would facilitate further exploration of EBLX/Ps. This is probably not limited to bornaviruses; there may be many hidden EVEs that are derived from small and/or rapidly evolving viral genes. Read-through transcription has also been observed for other mononegaviruses (26). Our search strategy can be applied to the detection of such EVEs. Additionally, a previous study identified virus-like insertions using a machine learning-based method, but the origins of insertions are still unclear (27). Our approach might also contribute to understanding the origins of those sequences.

Taken together, we identified EBLs that were undetectable by conventional BLAST searches. Our findings provide novel insights into ancient bornaviruses and co-evolution between bornaviruses and their hosts. Furthermore, our search strategy will contribute to paleovirological research.

## Materials and Methods

### Sanger sequencing of EBLs and flanking loci in M. fuliginosus

Genomic DNA was obtained from YubFKT1 (kidney epithelial cell line) (28) in a previous study (13). miEBLN/X/P-1, miEBLN/X/P-2, miEBLN/X/P-3, and miEBLX/P-4 and corresponding flanking loci in *M. fuliginosus* were amplified by PCR using the extracted DNA with Phusion Hot Start II High-Fidelity PCR Master Mix (Thermo Fisher Scientific). The PCR conditions are available upon request. The amplicons were purified using innuPREP PCRpure Lite Kit (Analytik Jena) and sequenced by Sanger dideoxy method at Eurofins Genomics. The primer sequences are available in S6 Table. The obtained sequences were deposited to DDBJ (accession numbers LC708263-LC708266).

### Cell lines

SuBK 12-08 cells (29) and human embryonic kidney 293T (HEK293T) cells were cultured in low-glucose Dulbecco’s Modified Eagle Medium (Nacalai Tesque) supplemented with 10% fetal bovine serum (Sigma-Aldrich) and 100 μg/mL penicillin-streptomycin (Nacalai Tesque). The cells were maintained at 37°C in a humidified chamber containing 5% CO_2_.

### Construction of miEBLX-4 and miEBLP-4 expressing plasmids

Expression vectors with C-terminal FLAG or Myc tag were first constructed. pcDNA3 was digested with EcoRI and XhoI, which was assembled with oligo DNA containing Myc or FLAG tag sequence using NEBuilder HiFi DNA Assembly Master Mix (New England Biolabs). The resultant plasmids were named pcDNA3-C-Myc or pcDNA3-C-FLAG. Then, miEBLX-4 and miEBLP-4 were amplified by PCR using the genomic DNA extracted from YubFKT1 with Phusion Hot Start II High-Fidelity PCR Master Mix, which were subsequently assembled with pcDNA3-C-Myc or pcDNA3-C-FLAG linearized with KpnI and EcoRI using NEBuilder. The oligo DNA sequences are available in S6 Table.

### Co-immunoprecipitation of miEBLX-4 with miEBLP-4

pcDNA3-miEBLX-4-myc and pcDNA3-miEBLP-4-FLAG plasmids (5 μg) were transfected into 293T cells on 10-cm dish using 20 μL of Avalanche-Everyday Transfection Reagent (EZ Biosystems) with Opti-MEM (Thermo Fisher Scientific). One day post-transfection, the cells were washed twice with ice-cold PBS and lysed using 500 μL IP buffer (150 mM NaCl, 20 mM Tris-HCl pH 7.4, 1 mM EDTA, 1% Triton X-100, 1× protease inhibitor cocktail [Nacalai Tesque]) for 30 min at 4°C with rotation.

In parallel, 500 μg SureBeads Protein G magnetic beads (Bio-Rad) were incubated with 3 μg mouse monoclonal anti-DDDDK antibody FLA-1 (M185-3L; MBL) in IP buffer for 1 hr at 4°C with rotation. The lysates were then centrifuged at 12000 × g for 10 min at 4°C, and then the supernatant were immunoprecipitated with the antibody-bead complex in IP buffer for 2 hr in 4°C with rotation. After incubation, the beads were washed thrice with ice-cold IP buffer and then transferred into new microtubes. The proteins were finally eluted from the beads by boiling in SDS sample buffer for 5 min at 95°C.

### Western blotting

Protein samples were separated in 12% polyacrylamide gel in Tris-glycine-SDS buffer, which were transferred to PVDF membrane (MERCK) at 10 V for 1 hr using Trans-Blot Turbo Transfer System (Bio-Rad). Then, the membrane was blocked with 5% skim milk in 0.1% Tween 20 in PBS (PBST) for 1 hr at room temperature (RT). The membrane was incubated with mouse monoclonal anti-Myc My3 (MBL; 1:5000) or anti-DDDDK FLA-1 antibody (MBL; 1:2000) diluted in 5% skim milk for 1 hr at RT, washed, and then incubated with horseradish peroxidase-conjugated donkey polyclonal anti-mouse IgG (715-035-150, Jackson ImmunoResearch; 1:2000) antibody diluted in 5% skim milk for 45 min at RT. The membrane was washed again, after which chemiluminescence reaction was conducted using Chemi-Lumi One (Nacalai Tesque) and was subsequently viewed using Amersham Imager 680 (GE Healthcare, IL, USA).

### BLAST search against bat genomes to detect EBLN/X/Ps and EBLX/Ps

BLASTn (14) search was conducted against the WGS database (15) of bat (Chiroptera, taxid: 9397) using the nucleotide sequence of miEBLN/X/P-1 in *M. natalensis* as a query on the BLAST web server (https://blast.ncbi.nlm.nih.gov/Blast.cgi). Changes applied to the default parameters were as follows: E-value threshold = 10^−10^, word size = 7, no filter for low complexity regions.

### TFASTX/Y search against vertebrate genomes to detect EBLX/Ps

Vertebrate genomes were downloaded using NCBI Datasets and were used as the database for TFASTX/Y searches. TFASTX/Y searches, included in the FASTA36 package (30), were performed against each species of the vertebrate genomes using the putative amino acid sequences miEBLX-4 and miEBLP-4 as queries with the ktup value option of 1. The obtained hits with E-values less than 10^−4^ were extracted, which were used for the downstream analyses.

### Identification of orthologous and syntenic empty loci

BLASTn search was conducted against the WGS database of bats (Chiroptera, taxid: 9397) or rodents (Rodentia, taxid: 9989) using the nucleotide sequences of EBL and its flanking regions as queries on the BLAST web server. Changes applied to the default parameters were as follows: E-value = 10^−20^, word size = 11. The BLAST hits which comprised of genomic sequences from different species were retrieved, and then aligned by MAFFT using the E-INS-i algorithm (31) in Geneious version 11.1.5. The nucleotide accessions and region of EBLs used for search queries are available in S7 Table.

### Annotation of EBLs and TSDs

TSD sequences were determined based on the alignment made in “*Identification of orthologous and syntenic empty loci*”. To identify transcription signal-like sequences, the nucleotide sequences of EBLs and extant orthobornaviral N/X/P or X/P transcripts were aligned by MAFFT using the E-INS-I algorithm. The transcription signal-like sequences and poly-A stretches were determined according to the resultant alignment.

To identify conserved functional domains of the miEBLX-4p and miEBLP-4p, the amino acid sequences of miEBLX-4p and miEBLP-4p in *M. fuliginosus* were aligned with extant orthobornaviral X and P proteins, respectively, by MAFFT using the E-INS-i algorithm. Based on the alignments, the conserved domains were manually analyzed.

The nucleotide accession of extant orthobornaviruses used in the analysis are available in S8 Table.

### mRNA sequencing analysis

Total RNA was extracted from SuBK12-08 cells using NucleoSpin RNA plus (MACHEREY-NAGEL) according to the manual. Sequencing library was constructed using TruSeq Stranded mRNA Library Prep Kit (Illumina). The libraries were sequenced by NextSeq 500 using NextSeq 500/550 Mid Output Kit v2.5 (300 Cycles).

The obtained reads were preprocessed by fastp 0.23.2 (32) with the default setting, which were mapped to the reference genome of *M. natalensis* (GCF_001595765.1) by HISAT2 version 2.2.1 (33) with the default setting. The mapped reads and splicing junctions were visualized using Integrated Genomics Viewer 2.13.0 (34).

The RNA-seq data were deposited to DDBJ Sequence Read Archive (DRA) under the accession number DRR403400.

### Pairwise similarity analysis

Amino acid sequences of the viral proteins of extant orthobornaviruses (S8 Table) were analyzed by MAFFT alignment tool using Sequence Demarcation Tool software version 1.2 (35). Minimum value for the percent identity scale was set to 0.1. The resulting matrices were set to 3-color mode and exported.

### In silico characterization of putative EBLX and EBLP proteins

To predict the protein secondary structure, amino acid sequences of miEBLX-4 and miEBLP-4 in *M. fuliginosus*, and extant orthobornaviral X and P proteins, were analyzed by JPred version 4 (36).

To determine the protein hydropathy, the same amino acid sequences were submitted to ProtScale (37) in ExPASy Server, using the amino acid scale developed by Kyte & Doolittle. The window sizes were set to 9 and 15 for X and P proteins, respectively.

## Acknowledgments

We are grateful to Ayato Takada for kindly providing SuBK 12-08 cell. We would like to thank Ken Maeda for kindly sharing YubFKT1. We are grateful to Takefumi Kondo and Yukari Sando for conducting the mRNA-sequencing. The animal silhouette images were downloaded under public domain access from https://commons.wikimedia.org/ and https://publicdomainvectors.org/.

This study was supported by JSPS KAKENHI grant numbers JP21H01199 (MH), 20H05682 (KT), JP21K19909 (KT), JSPS Core-to-Core Program JPJSCCA20190008 (KT), and MEXT KAKENHI grant number JP19H04833 (MH).

