## Supplementary Figure for "The hidden diversity of ancient bornaviral sequences from X and P genes in vertebrate genomes"

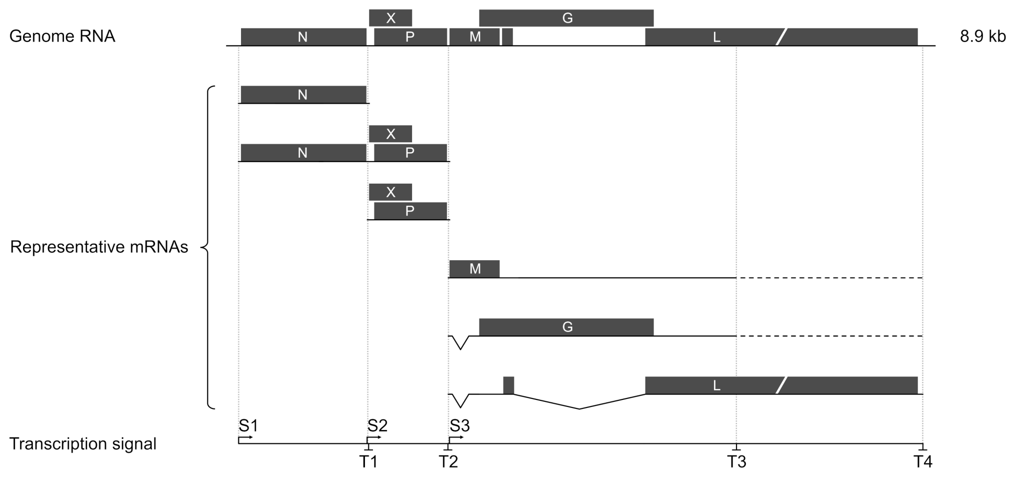


**Figure S1. Genome organization and transcripts of orthobornaviruses.** A schematic diagram of the genome organization and transcripts of orthobornaviruses.

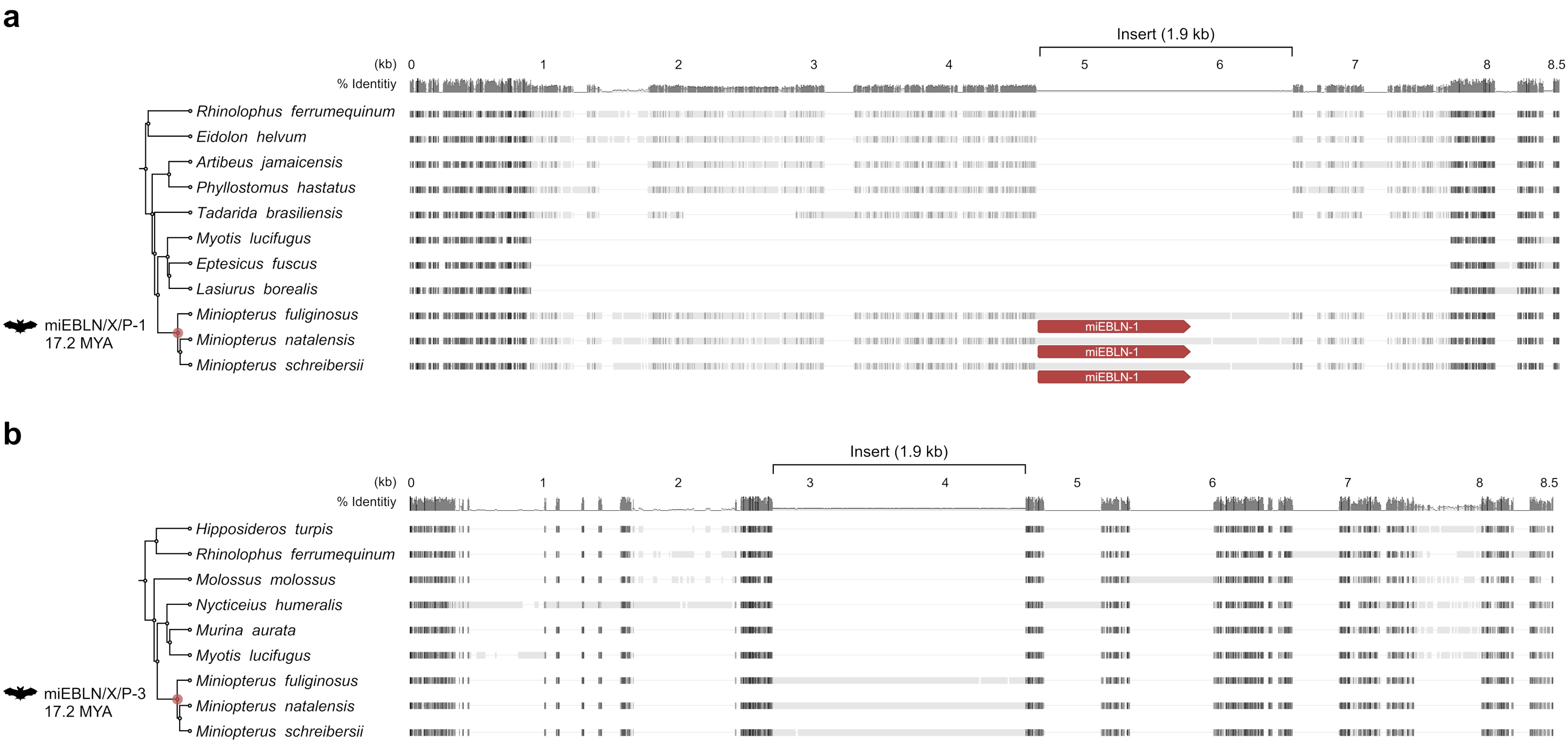


**Figure S2. Gene orthology analysis of EBLN/X/P-1 and EBLN/X/P-3.** Nucleotide sequence alignment of **(a)** miEBLN/X/P-1 and **(b)** miEBLN/X/P-3, and corresponding flanking loci are shown. Black and gray boxes indicate aligned regions. Horizontal lines show alignment gaps. Species phylogeny and deduced minimum integration age of each EBLN/X/P are indicated on the left based on TimeTree (1).

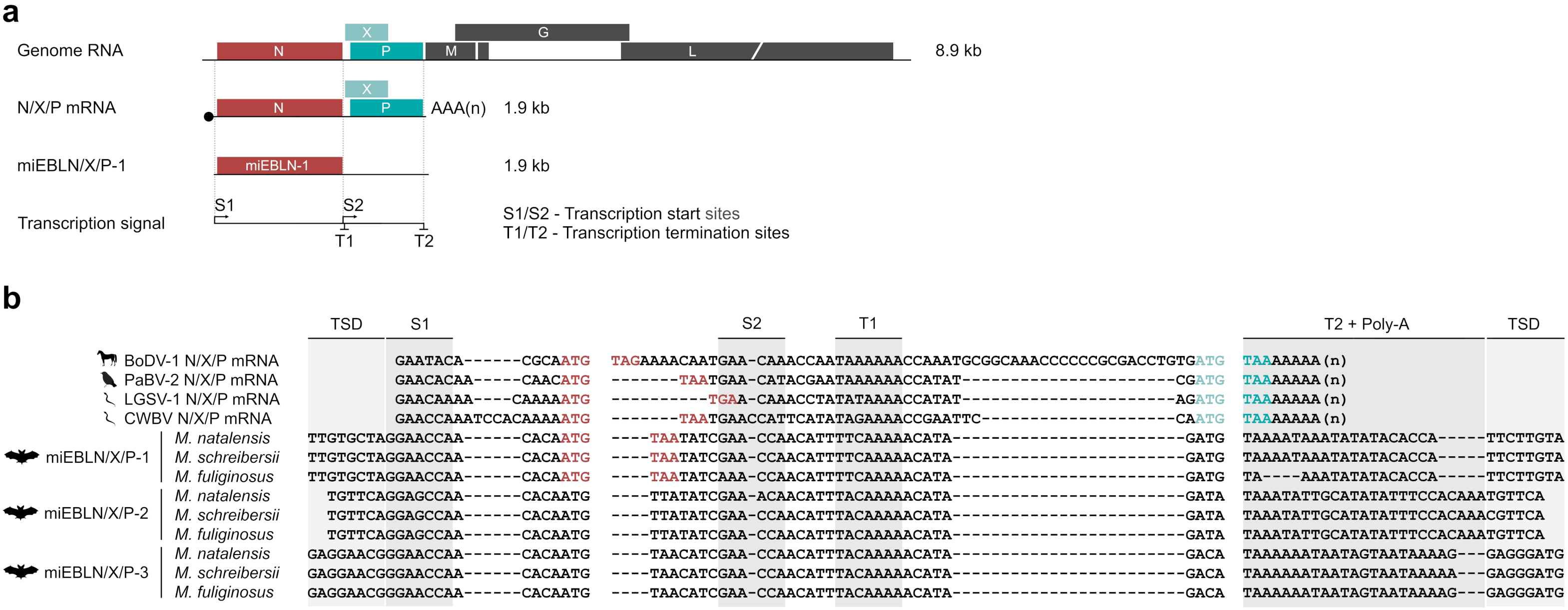


**Figure S3. Nucleotide sequence alignment of extant orthobornaviral N/X/P transcripts and EBLN/X/Ps. (a)** Schematic diagram of orthobornaviral genomic RNA, 1.9-kb N/X/P mRNA, and miEBLN/X/P-1. Transcription signals (S1, S2, T1, and T2) are shown. **(b)** Nucleotide sequence alignments of orthobornaviral N/X/P mRNA and miEBLN/X/P-1, -2, and -3. Red letters indicate the start and stop codons of the N gene. Light teal and dark teal letters indicate the start codon of X gene and the stop codon of P gene, respectively. TSD, target site duplication.

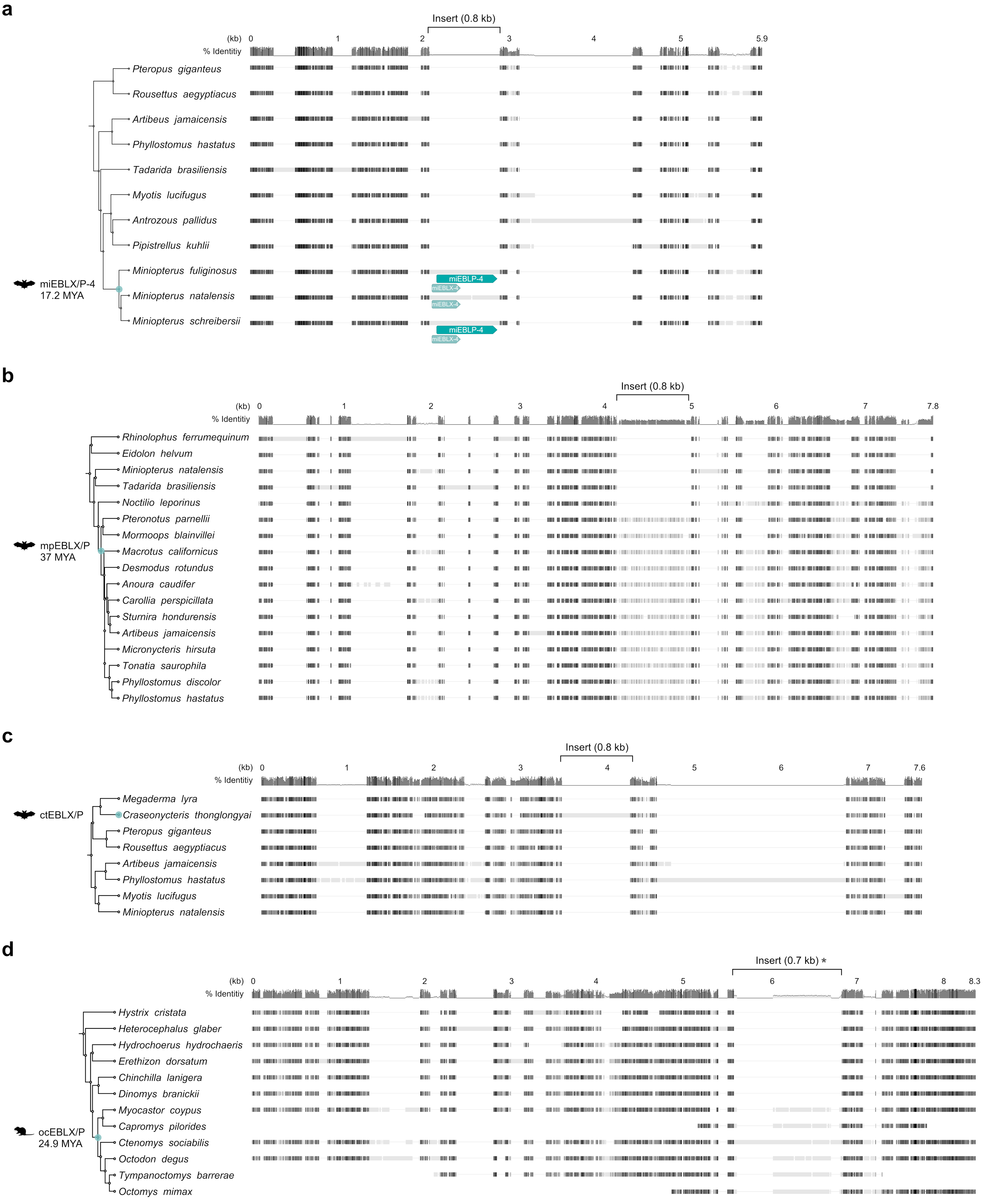


**Figure S4. Gene orthology analyses of EBLX/Ps.** Nucleotide sequence alignment of **(a)** miEBLX/P-4, **(b)** mpEBLX/P, **(c)** ctEBLX/P, and **(d)** ocEBLX/P, and corresponding flanking loci are shown. Black and gray boxes indicate aligned regions. Horizontal thin lines show alignment gaps. Species phylogeny and deduced minimum integration age of each EBLX/P are indicated on the left based on TimeTree (1). *ocEBLX/P in *C. sociabilis* contains insertions.

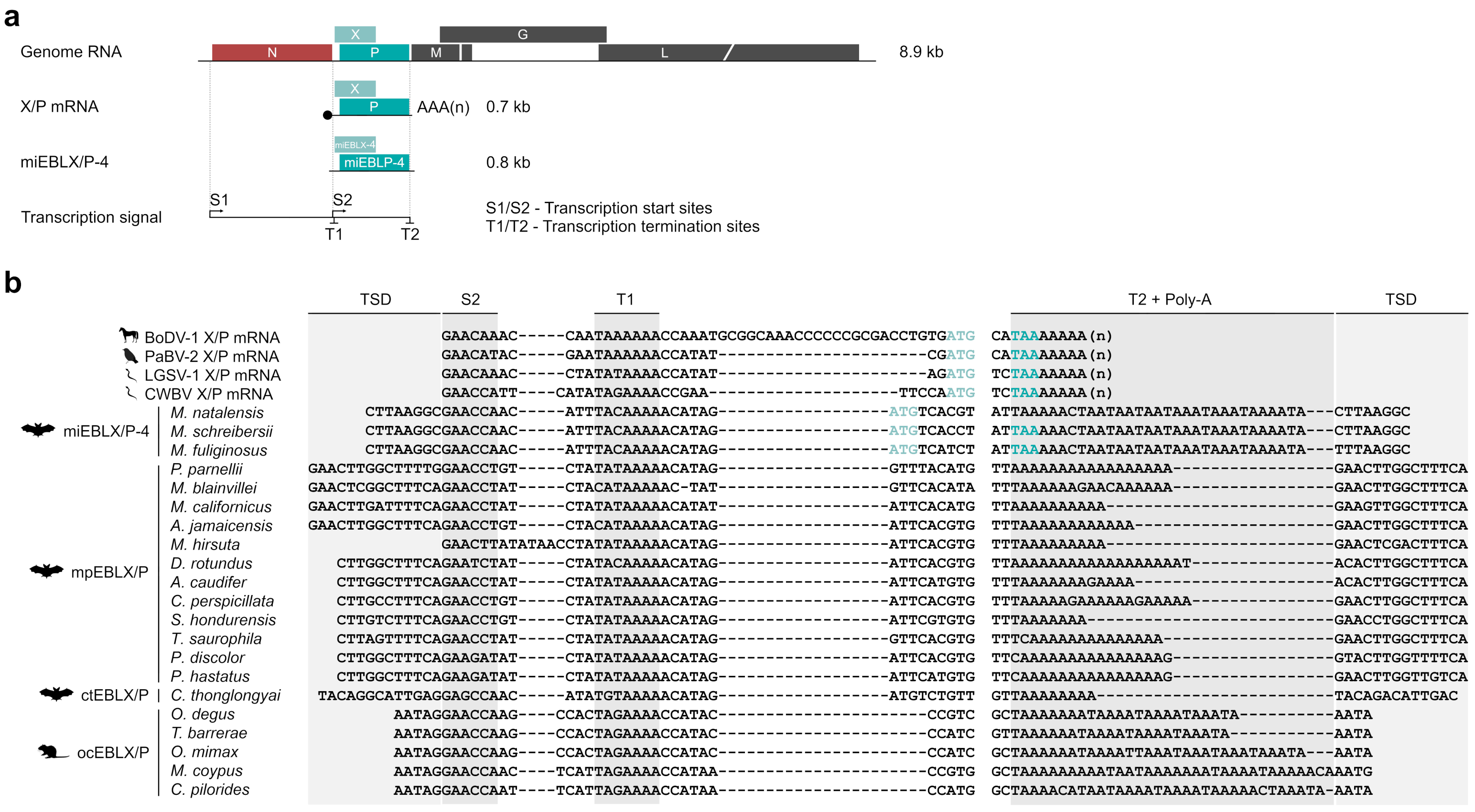


**Figure S5. Nucleotide sequence alignment of extant orthobornaviral X/P transcripts and EBLX/Ps. (a)** Schematic diagram of orthobornaviral genomic RNA, 0.7-kb X/P mRNA, and miEBLX/P-4. Transcription signals (S1, S2, T1, and T2) are shown. **(b)** Nucleotide sequence alignments of orthobornaviral X/P mRNA, and miEBLX/P-4, mpEBLX/P, ctEBLX/P, and ocEBLX/P. Light teal and dark teal letters indicate the start codon of X gene and the stop codon of P gene, respectively. TSD, target site duplication.

**a**

```
           1        10        20        30        40        50        60        70        80        90        100       109 (aa)

BoDV-1 X   MSSDLRLTLLELVRRLNGNATIESGRLPGGRRRSPDTTTGTIGVTKTTEDPKECIDPTGRPAPEGPQEEPLHDLRPRPANRKGAAIE
           ---HHHHHHHHHHHH--------------------------EEEE------------------------------------------

PaBV-2 X   MSTSFELTLLELIRRLNGTALEPSRVTRGGRRPADSTSNEVEVSKKEANTQGGPNCPRGETSRASQEESFHDLRSRTKDRKGAVIE
           --HHHHHHHHHHHHH-------------------------EEEEE-----------------------------------------

LGSV-1 X   MSADALLSIVELARRLDGDSKRQSSVSARRGPTVDSTSNQIEVTNKEEEESGQETAGVLSGRVGGQDVSKSGPNLRQRAAGNRKGSAVE
           --HHHHHHHHHHHHH-------------------------EE-----------------------------------------------

CWBV X     MNPNRFLCIVELLRRLNAKEAGDSCFPSGRGGNPDTTSEPEPLPNQEEGEVAKGSIDVEGGRTIEQDADRRGDYNLRSGGPSDRKGAAVN
           ----HHHHHHHHHHHH--------------------------------------------------------------------------

miEBLX-4   MSSEENKFLCMVKLLCRLHVGSRGTKLKGGGESGGDREAITSGDAKSKFQGKDSAQRERATPETHQDKAAPASFPDSSRQPKQECHSAGDCCNGKRYGEVDGSRGGNQA
           ------HHHHHHHHHHHHH------EE----------------------------------------------------------------------------------
```

H - α-helix
E - β-strand

**b**

```
           1        10        20        30        40        50        60        70        80        90        100       110       120       130       140       150       160       170       180       190       200       210       220       230 (aa)

BoDV-1 P   MATRPSSLVDSLEDEEDPQTLRRERSGSPRPRKIPRNALTQPVDQLLKDLRKNPSMISDPDQRTGREQLSNDELIKKLVTELAENSMIEAEEVRGTLGDISARIEAGFESLSALQVETIQTAQRCDHSDSIRILGENIKILDRSMKTMMETMKLMMEKVDLLYASTAVGTSAPMLPSHPAPPRIYPQLPSAPTADEWDIIP
           -----------------HHHHHHHH----------HHHHHHHHHHHHHHHH--------------------HHHHHHHHHHHH-----HHHHHHHHHHHHHHHHHHHHHHHHHHHHHHHHHHHHH---HHHHHHHHHHHHHHHHHHHHHHHHHHHHHHHHHHHH-------------------------------------

PaBV-2 P   MARPSSLVESLEEEDDPQTARRMRSRSPRRKRIPREALTVPVEKLLEQVKKNPSMISDQEPKTGREQLSNDDLIRQLITELADTNMIEAEGLKSSLEDIGSKLETGLESLSSLQIETLSAVQQSDYANSIKVLGENMRVLDRSIKTMNETMKMMMEKIDLLYSTMAIGNPTATMLPSHPGPSRIYPTLPTAPTAEDLDIIP
           ----------------HHHHHHHH----------HHHHHHHHHHHHHHHH--------------------HHHHHHHHHHHH-----HHHHHHHHHHHHHHHHHHHHHHHHHHHHHHHHHHH---HHHHHHHHHHHHHHHHHHHHHHHHHHHHHHHHHHHH---------------------------------------

LGSV-1 P   MATRRDNLVSPLEEDPPSTAHRTRSRSPIRRKKSLDKRLLGYSVEELVDKMSVNPGLIYDKEQLATGREALSSEELIKQLMGEIKTATEERKIDQEVIQMKFSHLERTLESYFESVVTTQVEILEKLGQLDYSGSIRQLGENMKILDKSLKAVTASVTLMTEKVDLLYGKMAVGTSNAPMIPSCPQPQSIYPKLPESTATAPALDIVF
           ------------------HHHHHHH-----------HHHHHHHHHHHHHHHH---------------HH----HHHHHHHHHHHHHH-----HHHHHHHHHHHHHHHHHHHHHHHHHHHHHHHHHHHHH-----HHHHHHHHHHHHHHHHHHHHHHHHHHHHHHHHHH-------------------------------------EE-

CWBV P     MPKKPETLVSPLEEEEIPTQPRNRSRSPIRKKGKLPKDLLTSRVEELSSKMRTEGVTTILDPGGPLTGKEQQSTSALIEELLQELRGEMEGQAIDRTELMSNIESIRVLSESLIGGQENILHKIELSDHAASIKQLGENVKILDRAIKAQGDVLTGLANKIDLIYARMAISDPKASMIASASGPSSLYPTLPTAPKPIDEIDIIF
           -------------------HHHHH------------HHHHHHHHHHHHHHH------EEE-------------HHHHHHHHHHHHHHH----HHHHHHH---HHHHHHHHHHHHHHHHHHHHHHHH---HHHHHHHHHHHHHHHHHHHHHHHHHHHHHHHHHHHHH---------------------------------------

miEBLP-4   MLGLEELSSKEEESQAEIERPSHQEMPSQSSKARTPPRGKEPPQRPTRTKRPQRVSRIPPGNLNKNVTQLVTAAMARDMEKLMDPEVETRLEERSTEELVQHVTAKNLTDQKVTSLETSINALAVFVGQVHEKQTEILSLIKMLPTKDDIKVLNKNVLQVASSVQQMSESINQITEKLNYVHARLAIQDPMPVMTPATAIVSSAPPASALSLYPVADLAVATSSYSVGDLVLD
           ----------------------------------------------------------------HHHHHHHHHHHHHHHH-------HHH-----HHHHHHHHHHH---H-HHHHHHHHHHHHHHHHHHHHH-HHHHHHHHHH------HHHHHHHHHHHHHHHHHHHHHHHHHHHHHHHHHHHHH-------------------------------HHHH--------EE--
```

**Figure S6. Secondary structure prediction of putative EBLX and EBLP proteins.** Predicted secondary structure of **(a)** extant orthobornaviral X and miEBLX-4 proteins, and **(b)** extant orthobornaviral P and miEBLP-4 proteins. H, α-helix. E, β-sheet.

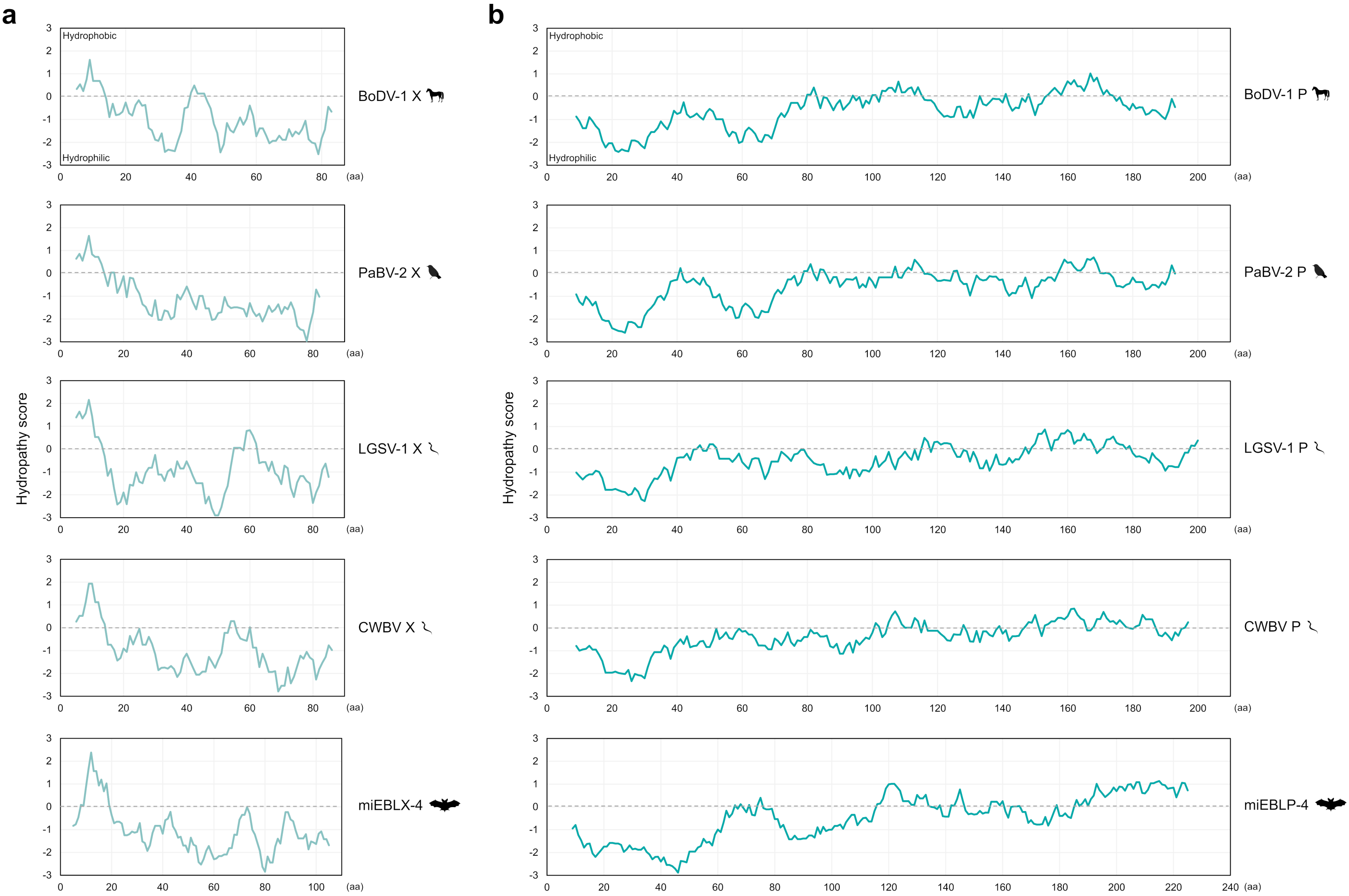


**Figure S7. Hydrophobicity of putative EBLX and EBLP proteins.** Hydropathy plot of **(a)** extant orthobornaviral X and miEBLX-4 proteins, and **(b)** extant orthobornaviral P and miEBLP-4 proteins.
